# Direction and modality of transcription changes caused by TAD boundary disruption in *Slc29a3/Unc5b* locus depends on tissue-specific epigenetic context

**DOI:** 10.1101/2024.08.16.608309

**Authors:** Paul Salnikov, Polina Belokopytova, Alexandra Yan, Emil Viesná, Alexey Korablev, Irina Serova, Varvara Lukyanchikova, Yana Stepanchuk, Nikita Torgunakov, Savelii Tikhomirov, Veniamin Fishman

## Abstract

Topologically associated domains (TADs) are believed to be involved in the regulation of gene expression. While the impact of TAD perturbations is usually studied in developmental genes with highly cell-type-specific expression patterns, this study examines genes with broad expression profiles divided by a strong insulatory boundary. We focused on mouse *Slc29a3/Unc5b* locus, which encompasses two distinct TADs. Our analysis demonstrates that deletions of CTCF binding sites within this locus lead to alterations in local chromatin architecture, disrupting existing loops and forming novel long-range interactions. We evaluated the transcription changes of *Unc5b, Slc29a3, Psap, Vsir, Cdh23*, and *Sgpl1* genes across various organs, finding that TAD boundary disruption results in variable transcriptional responses, where not only magnitude, but also direction of gene expression changes are tissue-specific. Current models of genome architecture, including enhancer competition and hijacking, only partially account for these transcriptional changes, indicating the need for further investigation into the mechanisms underlying TAD function and gene regulation.

## Introduction

In vertebrates, interphase chromatin is organized into Topologically Associated Domains (TADs) and sub-TAD loops by the interaction of the DNA-looping activity of the Cohesin complex and the DNA-binding insulatory factor CTCF [1]. Since this organization influences the spatial interactions of cis-regulatory elements, it is believed that TAD structure plays a crucial role in controlling gene expression. Despite intensive research, the role of TADs in transcription regulation remains controversial. There is a high conservation of TAD structure across species [2–5], and some experiments have shown that perturbations in TADs can have a significant impact on the expression of adjacent genes [6–8], leading to pathological phenotypes. However, other evidence indicates that TAD perturbations often have minor effects on gene expression[9,10] and do not always lead to phenotypic consequences. Therefore, interpreting TAD boundary mutations and predicting their effects on the magnitude and direction of expression changes in specific tissues remains challenging [11]. Previous studies discovered several mechanisms that explain the consequences of TAD boundary mutations. TADs limit the range of enhancer activity, and alterations in the evolutionarily shaped TADs structure can lead to the miswiring of regulatory elements and ectopic activation of genes, a phenomenon known as enhancer hijacking [12–14]. These improper enhancer-promoter interactions often result in a gene’s overexpression or ectopic expression[8,15]. In cases involving oncogenes or developmental genes, which require tight expression control, TAD boundary mutations can lead to cancer [16,17] and developmental disorders [6,15,18].

Cases where TAD structure is required to bring promoters and enhancers together and TADs disruption disconnect these genomic elements from each other, causing downregulation of gene expression are relatively less abundant [19]. However, there are some well studied examples of these mechanisms, particularly those involving developmental genes *C-MYC* [20,21], *SHH* [22,23] and *Pax3* [24].

Furthermore, the structural integrity of TADs influences gene regulation not only through enhancer and promoter interactions but also by affecting the distribution and maintenance of chromatin marks. TAD boundaries can block the propagation of histone modifications by chromatin remodelers, and losing their insulatory function can lead to the establishment of inappropriate chromatin domains [25]. This is presumably true for both active and repressive histone marks in vertebrates [26].

The effects of TAD perturbations are typically studied in developmental genes, which display highly cell-type specific expression signatures. Although the difference between developmental and housekeeping genes is not well-defined, studding genes silenced in most cell types may be biased because they are unresponsive to the effects of TAD boundary perturbations in the majority of epigenetic contexts. Conversely, there is substantially less data on the effects of TAD perturbations on genes, which are ubiquitously expressed across various cell types. It is assumed that the fusion of regulatory landscapes hosting multiple ubiquitously expressing genes could lead to the rewiring of regulatory elements, driven by competition between promoters for regulatory activity [27].

To further investigate this, in this study we focus on genes that have widespread activity profiles but are divided into two TADs by a strong insulatory boundary. To assess the regulatory effects of chromatin architecture in this context, we selected mouse *Slc29a3/Unc5b* locus, which contains two distinct TADs (Fig. 1A). The centromeric domain TAD contains *Slc29a3, Cdh23, Vsir,* and *Psap* genes, which are crucial for viability and are involved in a number of developmental processes. The telomeric domain contains genes *Unc5b* and *Sgpl1* placed nearby its borders, and *Unc5b* has super enhancer signatures within its first intron, active in several organs, particularly in cerebellum.

**Figure 1.**
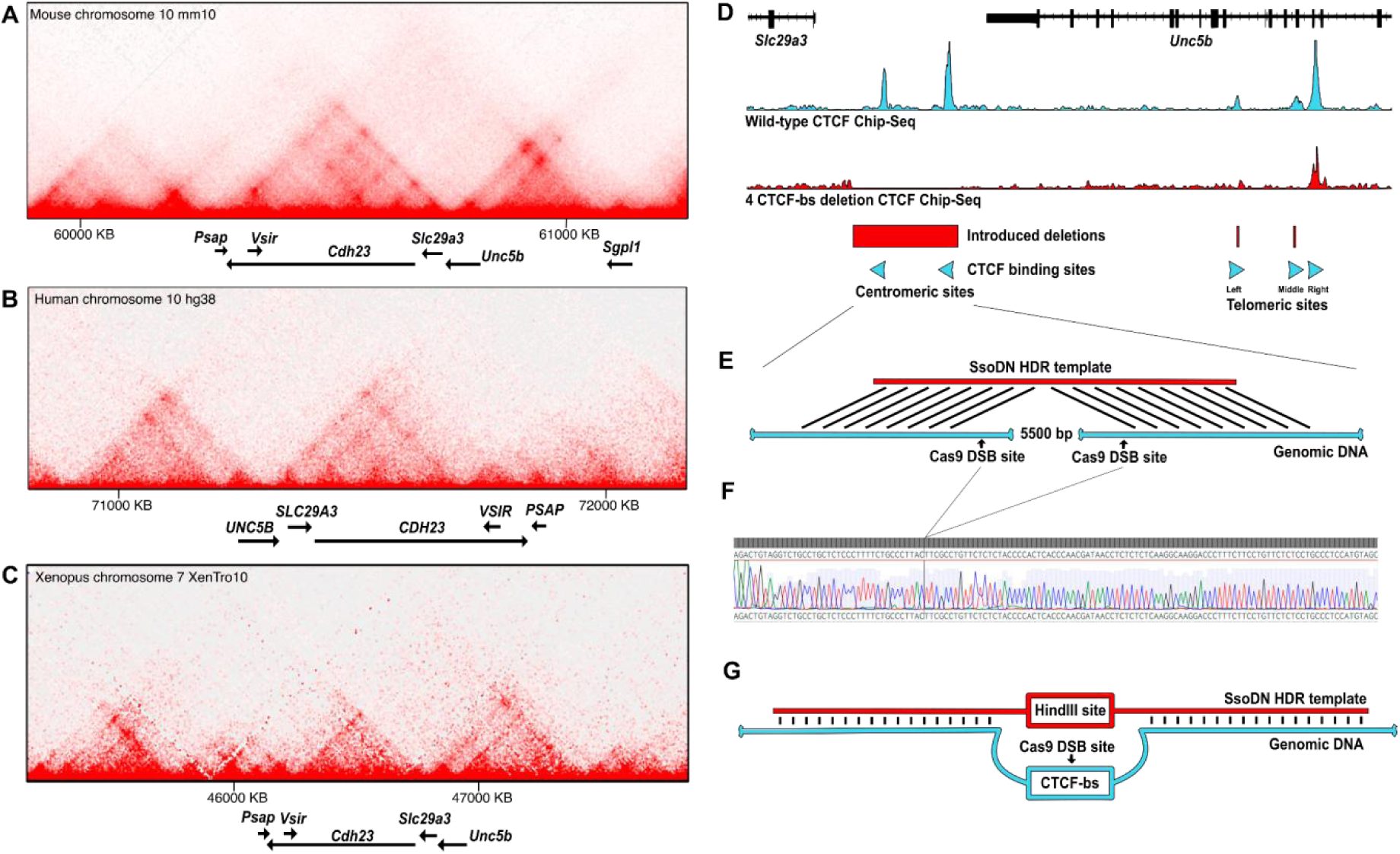
Spatial organization of the *Unc5b/Slc29a3* locus and its gene editing. A - Hi-C map and gene localisation of mouse *Unc5b/Slc29a3* locus. From [58] B - Hi-C map and gene localisation of human *UNC5B/SLC29A3* locus. From [59] C - Hi-C map and gene localisation of western clawed frog *Unc5b/Slc29a3* locus. From [60] D - CTCF-bs cluster at the mouse *Slc29a3/Unc5b* locus and CTCF ChIP-Seq tracks from mouse liver tissue (Wild-type and Mutant variant inherited from mouse #16). Mutations coordinates shown by red rectangles, CTCF-bs orientations shown by blue triangles. E - gene editing design employed for deleting the genomic region with two CTCF-bs. F - Sanger sequencing of the obtained deletion breakpoint regions. Obtained sequence aligned to the expected sequence of deletion. G - gene editing design for telomeric CTCF-binding sites knock-out.

Several genes located in the locus *Unc5b/Slc29a3* locus are clinically significant and play an important role in the development and functioning of organisms. *Unc5b* encodes the netrin-1 membrane receptor [28] that takes a part in axon [29] and vessel growth guidance [30,31], and also regulates proliferation, migration and apoptosis processes [32–35] and disruption of *Unc5b* expression is clinically significant marker in specific cancer subtypes [32,36,37]. In *Unc5b* knockout mouse models, the absence of *Unc5b* causes neurodevelopmental anomalies, embryonic growth arrest, and death during embryogenesis [31].

*Slc29a3* encodes a nucleoside transporter channel [38,39] that is important for cell homeostasis maintenance and autophagy regulation [40,41]. Its mutations are associated with hereditary scleroderma, hyperpigmentation, hypertrichosis, hypertrophy of internal organs, cardiovascular and musculoskeletal deformities [42] and immunodeficiency [43,44]; *Slc29a3* knockout mice exhibit splenic hyperplasia, hematopoietic dysfunction, and early death [40]. The *Cdh23* gene is associated with both syndromic and nonsyndromic genetic deafness [45,46]; absence of *Cdh23* in knockout mice leads to congenital deafness, pathologies of the organ of Corti [47–49], as well as various sensory and motor disorders [50]. Mutations in *Psap* can lead to severe conditions such as saposin deficiency syndrome with hepatosplenomegaly and atrophy of brain structures [51,52], and are also associated with an increased risk of parkinsonism [53]; *Psap* knockout mice show hypoactivity, demyelination, axonal degeneration, early death [54,55], and certain sensory-motor disorders comparable to the phenotypes observed in *Cdh23* knockouts [56,57].

This locus has a strong insulating boundary dividing it into two distinct TADs that is evolutionary conserved among vertebrates (Fig. 1A-C). In mice, this boundary is formed by five CTCF binding sites (CTCF-bs): two centromeric CTCF-bs have reverse-orientation and are located in the intergenic region without any functional marks ([mm10] chr10:60,757,172-60,757,204 and chr10:60,760,631-60,760,663); the remaining three sites are placed within *Unc5b* introns ([mm10] chr10:60,775,582-60,775,773, chr10:60,778,683-60,778,844, chr10:60,779,677-60,779,992) and have forward orientation (Fig. 1D).

We hypothesize that disruption of the TAD boundary in the *Slc29a3/Unc5b* locus may lead to dysregulation of these genes and shed light on the mechanisms that tie together the spatial genome organization and the establishing of transcription states in different tissues.

## Results

### Generation of genetically-modified mice

To investigate the role of chromatin architecture in the *Unc5b/Slc29a3* locus we generated two mouse lines with deleted TAD boundary forming CTCF-bs (Fig. 1D).

First, we deleted two centromeric sites by introducing ∼5kbp deletion (chr10:60,755,585-60,761,088, mm10). As they are placed in intergenic regions and do not overlap any known regulatory or coding elements, we apply the conventional CRISPR/Cas9 system to introduce two double-strand breaks, supplemented with Homology-Directed Repair (HDR) template, stimulating deletions formation (Fig. 1E). This results in the generation of animals with deletion of a 5 kb region containing both CTCF-bs. From 19 animals carrying the target modification, we picked a single male mouse for backcrosses and obtained the homozygous line. PCR-genotyping and Sanger sequencing confirmed that deletion occurred between target Cas9 cut sites and likely represent the HDR-product (Fig. 1F).

Further, we attempted to disrupt three telomeric CTCF-bs located within *Unc5b* introns. We refer to these telomeric binding sites as Left, Middle and Right sites, according to their arrangement on centromere to telomere axis (Fig. 1D). Close location to *Unc5b* gene elements prohibits usage of simple editing designs such as removing all three CTCF-bs via single extended deletion. Thus, we constructed three CRISPR/Cas9 sgRNAs that provided cleavage within the predicted CTCF motifs. To minimize the risk of unintended deletions between the cleavage sites [61], we engineered single-stranded oligodeoxynucleotides (ssODNs) to guide the repair process. This design ensures the replacement of the CTCF core motif with a HindIII restriction site. This modification serves two purposes: it disrupts CTCF binding and simplifies the genotyping of the modified animals. We based our approach on the assumption that during the repair process, a few nucleotides at the ends of the double-strand breaks (DSBs) are resected before new synthesis occurs using the ssODN as donor of homology (Fig. 1G).

In this experiment, we obtained 19 animals that were genotyped by PCR plus restriction fragment length polymorphism (PCR-RFLP) with enzymatic cleavage with the HindIII enzyme. This analysis showed that the expected mutation variant was obtained only in three mice (#1, 2, and 9) out of 19, and only in the one, middle CTCF site (Supplementary table 1). The observed mutation rate was significantly lower than anticipated, based on our previous studies[7]. Consequently, we hypothesized that the introduction of mutations through HDR was insufficiently effective. Therefore, we proceeded to investigate the presence of INDELs that might have resulted from other repair pathways. To achieve this, we analyzed PCR amplicons that encompass the CRISPR/Cas9 target sites using next-generation sequencing. The analysis, performed for each target site and for each animal individually, did not reveal any new HDR outcomes in addition to those detected by PCR-RFLP; however, we identified multiple alleles harboring INDELs at target regions (Fig. 2A-C). Based on the presence of short homology motifs flanking the INDELs, we assumed that many of the observed deletions were generated via microhomology end joining (MMEJ) repair pathway. We observed 24 MMEJ mutation occurrences (counting each variant from each animal), 23 NHEJ and only 4 HDR occurrences among 19 animals and three mutation points. We also found that mutation outcomes agreed well with inDelphi [62] software predictions.

**Figure 2.**
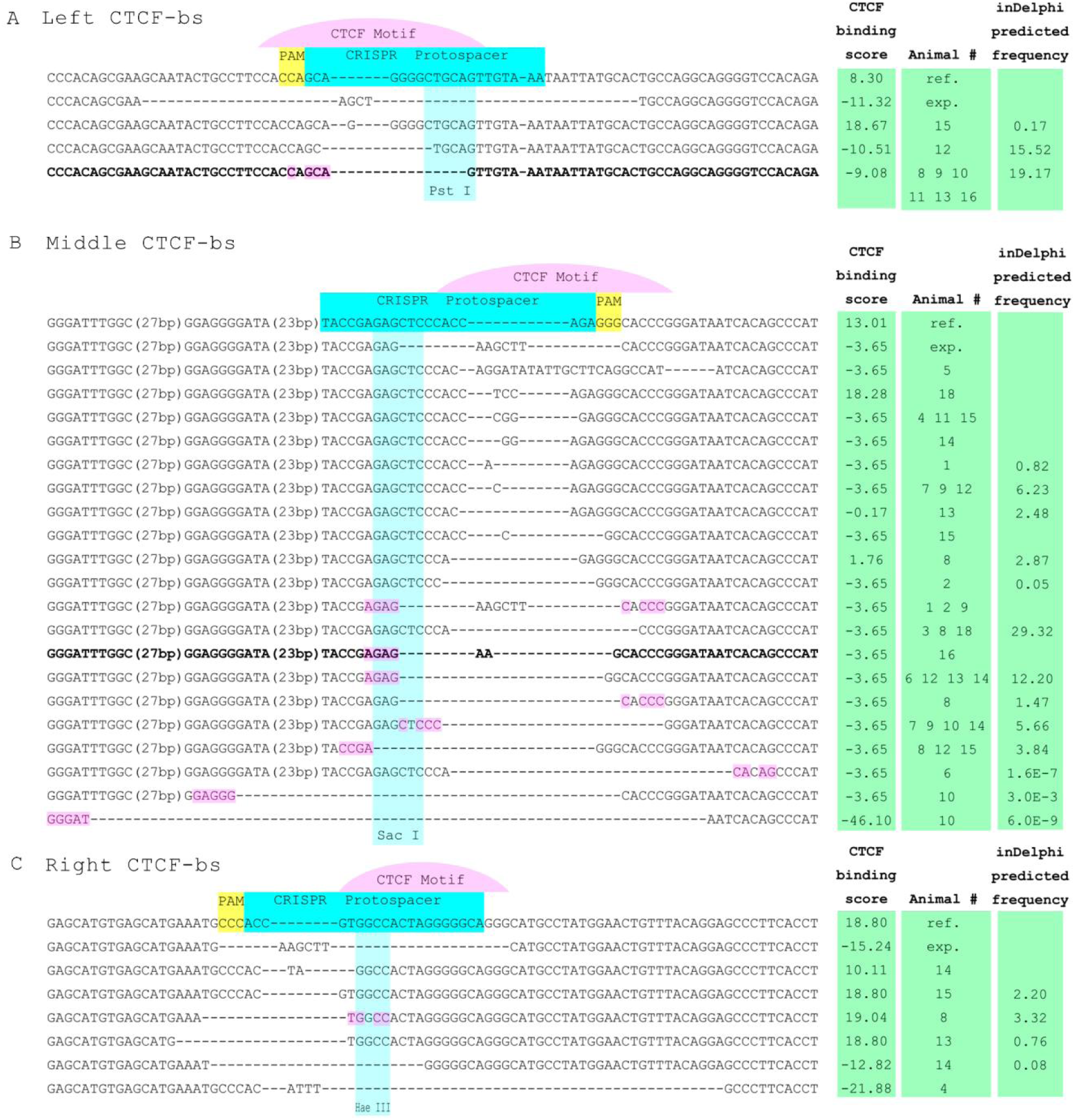
A-C - NGS genotyping results for Left (A), Middle (B) and Right (C) sites. ref. - wild-type allele sequence, exp. - expected sequence of HDR mediated mutation (equal to ssODN sequence). Potential microhomology motifs at deletion borders are highlighted by magenta. Variants chosen for homozygous line derivation (mouse #16) highlighted in bold.

To predict how mutations affect CTCF binding, we assessed the CTCF motif score of all generated sequences. We found that the majority of unexpected NHEJ and MMEJ deletions had low CTCF binding scores, comparable with binding scores predicted for intended HDR-mediated mutations. We chose the mouse #16, carrying a compound of 5,5-kbp deletion of intergenic CTCF-bs with Left and Middle mutations that have low CTCF binding scores, as most satisfying to derive a homozygous line.

Next, we confirmed that mutations of CTCF-bs affect CTCF binding. For this aim, we focused on liver tissue, where all five CTCF-bs in *Unc5b/Slc29a3* locus are bound by CTCF according to the public ENCODE data. Based on the ChIP-Seq experiment results on the homozygous animals liver, we confirmed the absence of CTCF binding for the mutated sites (Fig. 1D).

Thus, we obtained homozygous mouse line carrying deletions [mm10] chr10:60,755,585-60,761,088 (of two intergenic CTCF-bs, 5504 bp), chr10:60,775,689-60,775,698 (deletion of 9 bp at Left site of telomeric cluster), and chr10:60,778,725-60,778,738 (deletion of 13 bp with insertion of -AA- dinucleotide at Middle site of telomeric cluster), disrupting CTCF binding in this region.

### CTCF binding site deletions reconfigure local spatial contacts

To investigate the introduced mutations outcomes on spatial interactions of locus, we prepared capture Hi-C (cHi-C) libraries for cerebellum, kidney, and liver, using mice with four CTCF-sites deleted and wild-type controls (Fig. 3A-C).

**Figure 3.**
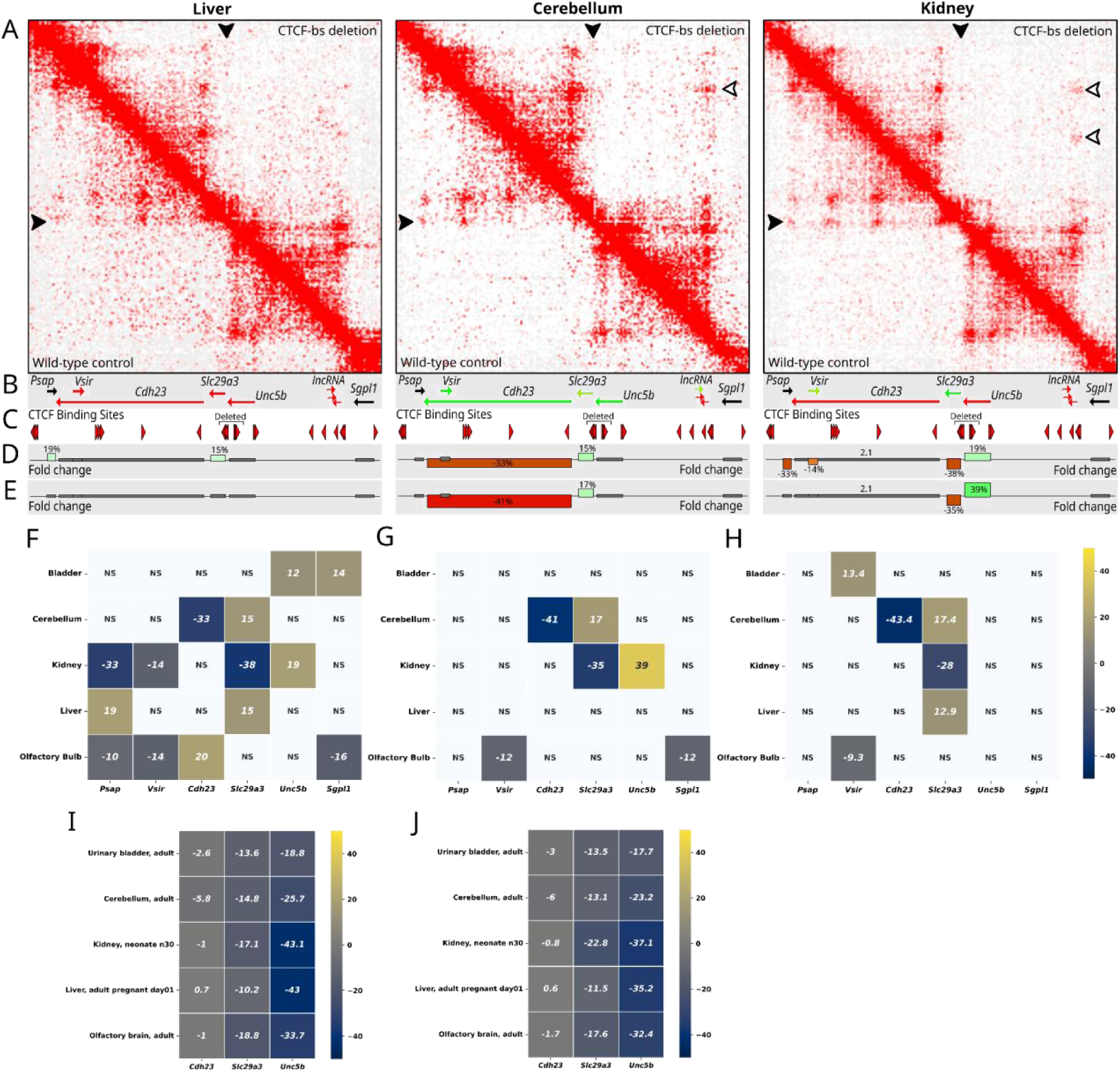
Alterations in Spatial Chromatin Architecture at the *Slc29a3/Unc5b* locus A - Hi-C Interaction Maps, displaying spatial chromatin interactions within the *Slc29a3/Unc5b* locus (chr10:60,103,000-61,356,000, mm10) in liver, cerebellum, and kidney tissues. Maps above the main diagonal represent mice with deletions of four CTCF-bs, while maps below the diagonal show data from wild-type mice. Arrows indicate chromatin loops differing between wild-type and CTCF-bs deletion conditions (refer to main text for details). B - Gene locations and their activity status that are indicated by a color scale from green (active expression) to red (repressed state). C - Location and orientation of CTCF binding sites. D, E - Bar plots representing changes in gene transcription within the locus for mice with deletions of two (D) or four (E) CTCF binding sites. Y-axis represent % of expression change relative to wild-type, X-axis reflects genomic coordinate, with bar width equal to gene length. F-J - Heatmap representations of expression changes. F-G - Mice with deletions of two (F) or four (G) CTCF-bs versus wild-type, NGS-based measurement. H - deletions of four CTCF-bs vs wild-type, digital PCR results. I,J - Enformer *in silico* predictions of the transcriptional effect of deletions of two (I) or four (J) CTCF-bs versus wild-type. Expression is shown as percentage of changes compared to wild-type, NS indicates non-significant results (FDR > 0.1).

In the wild-type state, the patterns of spatial contact were almost identical in all analyzed tissues. The centromeric TAD contains multiple loops. The three upstream loop anchors include: distal TAD boundary (located between the terminal end of the *Cdh23* gene and *Psap* gene), and two anchors within the *Cdh23* gene body. These three sites are forming the loops with downstream anchors, which are in the *Cdh23* promoter and within the TAD boundary located between *Slc29a3* and *Unc5b* genes. In the telomeric TAD, the loops are formed between the TAD boundary and *Unc5b* promoter and between two TAD boundaries. The distal boundary is placed near the cluster of long noncoding RNA genes between *Unc5b* and *Sgpl1* genes. It is noteworthy that almost all described loops are formed by convergent CTCF binding sites. This locus has only a single exception in this case: the CTCF-binding sites at the distal TAD boundary of the *Slc29a3* TAD align in the same direction as those at the anchors it interacts with.

The spatial maps of chromatin contacts in animals with CTCF-bs deletions revealed subtle changes at the subTAD level. We confirmed that loops formed by deleted CTCF-bs and directed into the *Slc29a3* TAD side were disturbed in all tissues (Fig. 3A, black-filled arrowheads). In wild-type animals, these loops connect the TAD boundary region with CTCF-bs in the *Cdh23* gene body. At the same time, loops directed towards the opposite domain were preserved.

The chromatin region from TAD boundary to *Cdh23* promoter, containing the *Slc29a3* gene, gained spatial interactions with the whole *Unc5b* TAD. This alteration gradually becomes more noticeable from liver to cerebellum tissues, coinciding with overall locus activity. It is apparent that the TAD boundary has shifted to the *Cdh23* promoter site, transferring *Slc29a3* gene from *Cdh23* and *Vsir* to the spatial proximity with the *Unc5b* gene.

In addition to disruption of some loops, we noted the formation of new, ectopic long-range interactions. These new loops occur between the inner CTCF-binding sites of the *Cdh23* gene and the distant border of the *Unc5b* TAD, an area containing the cluster of non-coding RNA genes, commonly silenced. Notably, these interactions cross the insulatory TAD boundary. Furthermore, the pattern of these new loops differs significantly between tissues (Fig. 3A, white-filled arrowhead).

Overall, our analysis shows that alterations of CTCF binding sites in the *Unc5b-Slc29a3* locus causes changes in local chromatin architecture, disruption of loops and formation of novel long-range interactions between gene promoters and regulatory sequences.

### Reorganization of local spatial contacts leads to changes in the transcriptional activity of nearby genes

Reorganization of spatial contacts caused by deletions of CTCF binding sites may result in alteration of gene expression in this locus. To test this hypothesis, we performed expression analysis in the obtained mice. Previous studies have suggested that the magnitude of gene expression changes caused by TAD boundary perturbation might be relatively small. To obtain precision necessary to detect small expression changes, we developed advanced methodology based on allelic imbalance measurement. Specifically, we assessed gene expression alterations by utilizing hybrid models from F1 offspring, produced by crossing genetically modified mice with CTCF-bs deletions maintained on C57BL/6 background with wild-type CAST mouse strains. These hybrids possess distinct alleles that differ in exonic SNPs, allowing allelic transcripts discrimination. Thus, effects of CTCF binding sites deletion can be studied in the same tissue samples by comparing expression of CAST and C57BL/6 alleles. This design largely protects us from many biases since alleles share trans-regulating factors, so the only factor responsible for the differences in allele expression is the cis-environment.

For transcripts quantification, we generated cDNA with UMI-barcoded, transcript-specific primers. Following NGS-analysis of the obtained products allows discrimination and quantification of alleles. Using this strategy, we profiled three groups of male F1 hybrids, obtained by crossing CAST females with 1) males from wild-type C57BL/6 mice, and 2) C57BL/6 mice with two centromeric or 3) four (two centromeric and two telomeric) CTCF-binding sites deleted at the *Slc29a3/Unc5b* locus.

Using this approach, we quantified the expression of six genes within the targeted locus (*Unc5b*, *Slc29a3*, *Psap*, *Vsir*, *Cdh23*, and *Sgpl1*) in five organs (bladder, cerebellum, kidney, liver and olfactory bulb) (Fig. 3D-G). We identified 20 cases of significant gene expression changes caused by CTCF-bs deletions among 60 analyzed cases (FDR=0.1). As we expected from previous studies, the alterations in gene transcription observed were modest, not surpassing a 50% change. Intriguingly, the nature of these changes showed a tissue-specific pattern with variation not just in magnitude but also in direction of changes. For example, *the Slc29a3* gene <u>decreases</u> its expression in the kidney, while <u>increases</u> expression levels in the cerebellum. To confirm the observed changes in gene expression levels, we performed digital PCR on the samples with four CTCF-bs deleted (Fig. 3H).

### Enformer fails to predict expression changes caused by CTCF mutations

Next, we aimed to compare our experimental data with predictions made by the novel deep learning model Enformer [63], which has been developed to predict gene expression from DNA sequences. Our objective was to evaluate the applicability of Enformer for studying the regulatory function of CTCF-binding sites. The authors of Enformer paper demonstrated that enhancer and insulator sequences contribute most significantly to the prediction accuracy.

We hypothesized that applying the Enformer model would enable us to predict gene expression changes resulting from CTCF-binding site (CTCF-bs) mutations across various mouse cell types.

We simulated CTCF-bs deletions and predicted gene expression changes for three genes: *Cdh23*, *Slc29a3*, and *Unc5b*. These predictions were then compared with experimentally observed gene expression changes, measured by RNA-seq, in five different organs. For this comparison, we selected the most relevant cell types from Enformer’s mouse targets and predicted gene expression changes for the three genes (Fig. 3I,J).

We predicted changes in gene expression for all three genes under two types of mutations: 2 CTCF-bs deletions and 4 CTCF-bs deletions. The differences in predicted gene expression changes between these two mutation types were moderate. Additionally, the predictions across different cell types were similar, indicating a lack of cell type specificity. To evaluate the predictive accuracy of the Enformer model, we performed Pearson’s chi-squared test to assess how well the model predicted the direction of gene expression changes among different genes and cell types. The test yielded a p-value of 0.24, indicating no significant match between the predicted and experimental data. Specifically, the Enformer model consistently predicted a decrease in gene expression across all genes and cell types, with only the magnitude of decrease varying among different genes.

Notably, recent studies [64–66] have also reported that Enformer does not always correctly predict the direction and magnitude of gene transcription changes in individuals. These researchers assessed Enformer’s predictive accuracy for specific genotypes using the RosMAP and Geuvadis databases, which include whole genome sequencing and gene transcriptional activity data from a single cell type for individuals in the 1000 Genomes Project. The Enformer model often failed to capture minor variations in gene expression between healthy individuals.

Based on our data and published findings, we concluded that the Enformer model is not suitable for predicting moderate changes in gene expression.

### Transcriptional effects rarely align with known mechanisms

We next aimed to fit observed in NGS-experiment differences in gene expression to the known models of interactions between promoters and regulatory elements. Since we observed tissue-specific patterns of gene expression changes, we assume that promoter-enhancer interactions in this locus are influenced by the local epigenetic context and are presumably determined by the gene’s transcription activity in the particular tissue.

Case I. Expression of genes separated by TAD boundary becomes more similar after CTCF-bs deletion. The pattern of expression changes of closely located Slc29a3 and Unc5b genes suggest that their cis-regulatory landscapes may interfere with each other after TAD boundary disruption. In the kidney, we detected almost 40% decrease of Slc29a3 expression. In the same tissue, we observe upregulation of Unc5b. Increase of Unc5b expression ranges from 20% in animals with 2 CTCF-bs deleted to 40% in animals with 4 CTCF-bs detected. Notably, in this organ Slc29a3 is normally activated, while the Unc5b is repressed. Thus, we assume that when the insulator boundary between these genes is disturbed, they are influenced by the regulatory elements of each other. As a result, the expression of these genes is “averaging”.

Case II. Enhancer hijacking causes upregulation of low expressed genes. We detected approximately 15% increase in Slc29a3 expression in the cerebellum, where the expression of the Unc5b gene is significantly higher compared to its average level across mouse tissues. This suggests that Slc29a3 expression is this tissue is influenced by active cis-factors, for example Unc5b active super enhancer signatures within its first intron according to the ENCODE database. The Slc29a3 gene, by entering into spatial interaction with the Unc5b gene, also interacts with its chromatin environment, which explains the increase in its activity. In contrast to the aforementioned expression “averaging” observed in the kidney, there is no detectable decrease in the expression of the Unc5b gene, which can be attributed to the specific mode of its enhancer action or the small scales of these changes. In addition to the two cases described above, there are many other examples of gene expression changes that do not fit well into known mechanisms of gene regulation.

In the liver, both Unc5b and Slc29a3 genes are repressed; however, we observe weak Slc29a3 expression increase upon boundary disruption. Increased Slc29a3 expression can not be explained by interaction of the Slc29a3 gene with regulatory elements from the Unc5b spatial environment, because there are no candidate elements based on available ChIP-seq data, and it is not clear why if these elements exist in the Unc5b environment they do not cause active expression of the Unc5b gene.

In cerebellum, we observed significant reduction of the Cdh23 expression. There are two known mechanisms of gene silencing caused by TAD boundary disruption: disconnection of promoter from regulatory element and spreading of heterochromatin. We did not find any obvious candidate element with enhancer signature that changes contact frequency with Cdh23 promoter upon TAD disruption.

As shown by Hi-C experiments, the disruption of TAD boundary led to emergence of ectopic long-range interactions between the Cdh23 gene body and Unc5b TAD distal boundary. Noteworthy that these loops are observed only for tissues where Cdh23 is expressed (for cerebellum and kidney, but not for liver). We can speculate that these newfound interactions are linked with the spreading of repressive chromatin states, responsible for downregulation of the Cdh23 gene in cerebellum. However, we did not find any repressive chromatin marks located within anchors of the new Cdh23 loop.

In our experiment, the Psap gene exhibited the most intriguing dynamics. Despite being the furthest from the mutated TAD boundary region, this gene is downregulated in the kidney of animals with two CTCF-bs deleted. In contrast, it is expressed normally on the allele with four deleted CTCF binding sites. It is noteworthy that its pattern of change in the animals with two CTCF binding sites deleted closely resembles that of the Slc29a3 gene.

Psap and Slc29a3 genes are normally connected by the loop structure. This suggests that the parallel changes observed in these genes could be explained by their mutual influence through three-dimensional contact. Therefore, alterations in the epigenetic state of one gene within this hub (in our case, Slc29a3, which is influenced by Unc5b over the weakened TAD boundary) can be transmitted to other genes. However, when the TAD boundary is completely disrupted, severing the contact between Psap and Slc29a3, Psap expression level returns to normal.

We can also hypothesize that such congruent variation mediated by spatial looping may involve the Vsir and Sgpl1 genes in the kidney, liver, and olfactory bulb. Although the changes in these genes do not clearly follow the observed pattern, this could be due to an insufficient sample volume or interference from other effects. Finally, the changes observed in Unc5b and Sgpl1 in the bladder, and Cdh23 in the olfactory bulb, are not easily explained. In general, changes in gene expression tend to be opposite to their baseline activity level in a given tissue. Thus, in cases where expression levels change, active genes typically decrease their expression, while repressed genes show an increase.

## Discussion

Here we obtained two mouse lines carrying deletions of two or four CTCF-bs between Slc29a3 and Unc5b genes. We have shown that disruption of the TAD boundary, formed by these CTCF-bs, led to reorganization of spatial contacts with fusion of the Slc29a3 gene region to the Unc5b-containing domain and establishing new inter-TAD loops which have unknown mechanisms of formation. This reorganization has a tissue-specific manner, manifesting more in actively transcribed states of locus.

During line generation, we used a multiplex modification of adjacent CTCF-binding sites by CRISPR/Cas9 gene editing. Our strategy utilized ssODNs as HDR templates due to their predictable mutagenesis pattern [67]. However, this strategy proved to be ineffective because it was outcompeted by the MMEJ DNA repair pathway. Nevertheless, we were able to obtain allele carrying MMEJ-induced mutations disrupting two CTCF-bs simultaneously. Thus, we do not recommend use of ssODN-facilitated mutagenesis in murine zygotes to create CTCF motif-disrupting INDELs. Instead, MMEJ pathway showed a well-predictable specter of generated mutations that could be exploited for such purposes, even in multiplexed manner.

We demonstrated that the deletions we obtained disrupt the TAD boundary and alter the inner loop structure of the Slc29a3 TAD. In the organs analyzed, there was no fusion of domains; however, the boundary shifted to the inner CTCF-binding site in the promoter of the Cdh23 gene. This shift transferred the DNA region containing Slc29a3 into the new cis-environment of the Unc5b TAD. Additionally, we observed the establishment of new long-range inter-TAD spatial contacts, the appearance of which cannot be readily explained by any currently known mechanisms.

We developed a sequencing-based method allowing us to precisely quantify gene expression alterations caused by TAD reorganization. We showed that this quantification strategy is well commutable with such a precise method as digital PCR, and therefore could be applied for assessment of any other cis-acting mutation consequences.

We revealed a wide range of gene expression alterations, despite that, as we expected, expression modulation has low amplitude. These results agreed with other experiments even with ones that investigated cases of developmental genes[8,10]. Such a low impact on the molecular phenotype challenges the conservation of CTCF-bs sites and TAD structure. This discrepancy still remains a major obstacle for the whole theory of TAD function, and needs more detailed investigations to be released.

Revealed pattern of changes proved to be tissue-specific not only in terms of the amplitude of changes but also in the direction of these changes. Although it seems clear that the changes caused by TAD boundary disruption are conditioned by the epigenetic state of the locus, it is still hard to predict. Neither empirical models - such as enhancer hijacking model - nor statistical models such as Enformer can explain the full spectrum of the observed expression alterations. These results highlight the need for new tools for non-coding genetic variants of interpretation, required by medical and evolutionary genomics [11,68].

The hypothesis derived from our data is that actively transcribed genes tend to lose their level of expression, whereas repressed genes, oppositely, often gain new enhancer interactions when regions with different chromatin states are merged together. This effect was observed in our experiment for interaction of Slc29a3 and Unc5b genes that tend to averaging of its expression levels after losing insulation between them, and for Cdh23 gene that, being normally highly activated in cerebellum, lost its expression level upon TAD structure reorganization. Thus, depending on locus’ epigenetic state and differentiation trajectory, it is expected that TAD boundary lesion can have tissue-specific effect on gene expression varying not only by the amplitude, but, more intriguing, by the direction of alteration.

An intriguing discovery is that the gradual deletions of boundary-forming CTCF-bs do not consistently result in a proportional enhancement of the effect. Specifically, in the cases involving Psap and Vsir, partial disruption of the boundary had a more significant effect than its complete disappearance. Additionally, we observed that the effects on Slc29a3 did not escalate in tandem with the progression of boundary disruption. This indicates that deletions involving two and four CTCF-bs may impact different TAD boundary functions rather than simply exacerbating a single effect. Moreover, this observation leads to the fascinating hypothesis that bidirectional TAD boundaries operate under a different functional logic compared to those with only co-directed CTCF-bs.

In summary, here we demonstrated that the impact of TAD boundary disruption gene transcription regulation is highly tissue-specific, with their magnitude and direction varying from tissue to tissue, making it challenging to predict consequences of TAD boundary mutations. This indicates a complex interaction between TAD structure and epigenetic regulation during the cell differentiation process in establishing gene expression patterns. Additionally, many of the observed effects do not align with any known mechanisms, prompting new questions for further investigation.

## Materials and methods

### Mouse lines and microinjections

We used the C57BL/6 mouse line as a basis for derivation of mutant mice (obtaining zygotes and backcrossing). We used pseudopregnant female CD-1 mice for transplantation of microinjected zygotes. Cytoplasmic microinjection of zygotes was performed using standard techniques that are widely used in transgenesis [69,70]. Food and water were available for animals ad libitum.

All animal procedures were approved by the Ethics Committee of the institute of Cytology and Genetics (protocol #65, issued October, 09, 2020). Animals were obtained and handled in the SPF Animal Facilities of ICG SB RAS.

### CRISPR/Cas9 sgRNA construction

We designed CRISPR sgRNAs for desired regions using web-tool «Benchling» (https://benchling.com). Then we simulated desired mutation sequences and designed ssODN for it with 60bp homology arms and primers for genotyping (Table 1). In the ssODN sequence for Left, Middle and Right loci we introduced HindIII recognition site for genotyping convenience. DNA templates for sgRNA synthesis were obtained via PCR using oligonucleotides containing T7 promoter, guide sequence and sgRNA scaffold. PCR products were used for an in vitro transcription (MEGAshortscript™ T7 Transcription Kit, Ambion). Obtained RNA was purified on MEGAclear™ Transcription Clean-Up Kit (Ambion) columns, and mixed with spCas9 mRNA (GeneArt™ CRISPR Nuclease mRNA, Thermo, USA) in ratios 8,2 pmol each sgRNA and 16,4 pmol Cas9 mRNA. ssODNs were purified on MEGAclear™ Transcription Clean-Up Kit (Ambion) columns and mixed in 250 ng/ul final concentration. RNA mix and ssODN were mixed immediately before microinjection.

### Genotyping

Animal genotyping was performed using three week old animals. DNA from the tail tip tissue was purified using the Phenol-Chloroform method and amplified by standard PCR protocol using corresponding primers (Supplementary table 2).

Deletion chr10:60,755,585-60,761,088 was detected using primers UNC5B-M1-F and UNC5B-R1-R (product length 300 bp). Wild type allele was detected using PCR from primer pairs UNC5B-M1-F and UNC5B-M1-R (475 bp); UNC5B-R1-F and UNC5B-R1-R (450 bp). Inversions and duplications of region chr10:60,755,585-60,761,088 were accessed by UNC5B-M1-F and UNC5B-R1-F; UNC5B-M1-R and UNC5B-R1-R; UNC5B-M1-R and UNC5B-R1-F primer pairs.

For detection of mutations in Left, Middle and Right loci primers UNC5B-L2-F and UNC5B-L2-R (461 bp); UNC5B-M2-F and UNC5B-M2-R (276 bp); UNC5B-R2-F and UNC5B-R2-R (324 bp) were used. PCR products were digested by HindIII restriction enzyme to detect HDR outcomes, or PstI (Left), SacI (Middle), or HaeIII (Right) to detect INDELs. Products of these reactions were analyzed by electrophoresis in 2% agarose gel.

### NGS libraries construction for genotyping

We designed primer pairs for Left, Middle and Right regions so that the PCR product is 250 bp length. For each region we purchased four forward and five reverse barcoded primers, so combination of them allowed us to multiplex NGS sequencing of 20 mice. PCR products from each region and each F0 mouse DNA sample were mixed and prepared for sequencing using Kapa Hyper Prep Kit (Roche, #KK8504) and a KAPA Single-Indexed Adapter Set A, (Roche, #KK8701) according to the manufacturer protocol without post-ligation PCR to prevent formation of PCR chimeras. Library was sequenced in BGI and we obtained 25.5 mln 150 bp length paired reads.

We used cutadapt to remove adapters. Reads were mapped with BWA MEM utility using default configuration. Then demultiplexed we assumed that bases on specific positions in mapped reads are mouse-specific primer barcodes. All genomic variants present in reads, regardless of their frequency in mapped NGS data, were documented and scored according to the number of reads supporting the variant. We visualized the histogram of variant frequencies and defined an empirical threshold, thus filtering out all variants supported by less than 2000 reads. The resulting set of genomic variants was thus compiled and presented in the work.

Predictions of mutation variant frequency were obtained using inDelphi online software (https://indelphi.giffordlab.mit.edu). Predictions of CTCF-binding score of mutation variants were obtained using CTCFBSDB 2.0 *in silico* CTCF-sb prediction tool (https://insulatordb.uthsc.edu).

### ChIP-Seq

Freshly dissected liver samples (about one gram) were grinded with razor, homogenized in Dounce homogenizer in 1 ml of 3% formaldehyde, then samples were transferred in 50 ml tube and volume of 3% was adjusted to 40 ml. After 10 min incubation at RT the fixator was quenched by addition of glycine to 125 mM final, incubated 10 min and washed twice by ice-cold PBS, then snap-freezed and stored at −80°C. Frozen samples were thawed and lysed for 30minutes in a lysis buffer (10 mM Tris-HCl, 1 mM EDTA, 1% Triton X-100, 0.1% sodium deoxycholate, protease inhibitors) supplemented with 0.5% SDS. Chromatin was shared using a Bandelin Sonopulse sonicator (70% power within 11 cycles of 30/90sec ON/OFF). 20-40 μg of fixed chromatin were used per one ChIP. Before incubation with specific antibodies, chromatin was diluted with a lysis buffer up to 0.1-0.2% SDS and pre-cleared by incubation with Protein A magnetic beads (New England Biolabs, S1425S) for 2h at 4C with slow rotation. During this time, another aliquot of Protein A magnetic beads was washed in PBS, combined with 5 μg target antibodies, and incubated for 2h at 4C with slow rotation. Beads were removed and pre-cleared chromatin was immunoprecipitated with antibody/magnetic beads complexes and incubated overnight at 4C with slow rotation. Next day, the beads were thoroughly washed in a series of buffers (Buffer 1: 10 mM Tris-HCl, 1 mM EDTA, 1% Triton X-100, 0.1% SDS, 0.1% sodium deoxycholate, protease inhibitors; Buffer 2: 500 mM NaCl, 10 mM Tris-HCl, 1 mM EDTA, 1% Triton X-100, 0.1% SDS, 0.1% sodium deoxycholate, protease inhibitors; Buffer 3: 0.25 M LiCl, 10 mM Tris-HCl, 1 mM EDTA, 0.5% NP-40, 0.5% sodium deoxycholate; Buffer TE/Triton: 10 mM Tris-HCl, 1 mM EDTA, 1% Triton X-100; TE buffer: 10 mM Tris-HCl, 1 mM EDTA). Cross-links were removed and DNA was eluted in 100 μL elution buffer (10 mM Tris-HCl, 1 mM EDTA, 1% SDS) by incubation at 65C for 14 hours. After treatment with RNAse A (New England Biolabs, T3018) and Proteinase K (New England Biolabs, P8107S), magnetic beads were removed. DNA was extracted using ChIP DNA Clean & Concentrator columns (Zymo Research, D5205). ChIP-seq libraries were prepared for sequencing using Kapa Hyper Prep Kit (Roche, #KK8504) and a KAPA Single-Indexed Adapter Set A, (Roche, #KK8701). Libraries were sequenced on DNBSEQ sequencing platform. ChIP-Seq data were processed by the standard ENCODE pipeline (https://github.com/kundajelab/chipseq_pipeline).

### cHi-C

Capture Hi-C library preparation was performed according to the original Hi-C 2.0 protocol[71] with minor adaptations for tissue samples proceeding:

Tissue samples were manually minced using a sharp blade and then suspended in 2 ml of PBS. The mixture was homogenized using a glass homogenizer for 20 strokes on ice. Formaldehyde was added to the final concentration of 2%, and the mixture was incubated for 10 minutes at room temperature with rotation. Glycine was then added to a final concentration of 250 mM and incubated 10 minutes under the same conditions. The samples were centrifuged at 1000g for 5 minutes, followed by two washes with cold PBS, after which the supernatant was discarded. The pellet was quickly frozen in liquid nitrogen or directly resuspended in a cold lysis buffer (10 mM Tris, 10 mM NaCl, 0.2% Igepal CA-630). Homogenization was performed using a syringe, and the lysate was filtered through muslin. The homogenate was incubated for 30 minutes on ice, centrifuged at 600g for 5 minutes, and then washed twice with cold lysis buffer and once with NeBuf 3.1 containing 0.3% SDS. The pellet was resuspended in NeBuf 3.1 with 0.1% SDS and incubated for 30 minutes at 37°C, followed by the addition of 200 µl of 1.5% Triton X-100 and further incubation under the same conditions. DpnII enzyme (5 µl) was added, and the mixture was digested overnight at 37°C. After a 20-minute incubation at 65°C, the mixture was centrifuged to remove the supernatant. The DNA was then prepared for end repair by resuspension in 150 µl of DNA end repair mix (50 µM dGTP, 50 µM dATP, 50 µM dTTP, 50 µM dCTP-15bio, 1X NEBuffer 2.1, 25U PolII Klenow fragment), and incubated for 4 hours at 23°C. Ligation was performed by adding 1 ml of Ligation mix (1X T4 Ligase Buffer, 1% Triton X100, 5% PEG, 100 µg/ml BSA, 1 mM ATP, 4000U T4 Ligase) and incubating overnight at 16°C. The procedure was then continued as specified in the original protocol. Sequencing libraries were prepared using the KAPA Hyper Prep kit. The hybridization of libraries with RNA probes was performed according to the myBaits Manual v4.01 (Arbor Biosciences). Enrichment probes were designed over the region chr10:60,103,000-61,356,000, mm10.

### dPCR

The sample sizes in each group and organ combination varied due to differences in sample preparation quality and the availability of experimental mice. However, each group in this experiment consistently included at least five samples.

To perform digital PCR with fluorescent probes, the QIAcuity® Probe PCR Kit (Qiagen) was used. According to the protocol, for one 12 µl reaction, the following were mixed: 3 µl of 4x Probe PCR Master Mix, 1 µl of each corresponding primer (10 µM) and probe (10 µM), and 0.5 µl of cDNA sample. The reaction mix was transferred to reaction nanoplates and sealed with specialized film (from the QIAcuity Nanoplate Seal, Qiagen). The plates were placed in the QIAcuity One device (with a 5-channel sensor), and the amplification reaction was performed as follows: reaction initiation at 95°C for 2 minutes; 35 cycles at 95°C for 15 seconds, 62°C for 30 seconds.

### Allele-specific NGS-based expression measurement

The sample sizes in each group and organ combination varied due to differences in sample preparation quality and the availability of experimental mice. However, the control group consistently included at least five samples, while both groups of mice carrying two or four CTCF-binding site deletions included at least six samples each.

To construct NGS libraries to get information about comparative allele expression levels, we first conduct reverse transcription from gene-specific primers, containing UMI and sequencing adapter part at its 5’-end, using RNAScribe reverse transcription kit (Biolabmix, Russia) using the manufacturer’s protocol. Next, we amplified cDNA using gene-specific second strand synthesizing primer and primer annealing at the sequencing adapter part of the reverse transcription primer. These amplicons were indexed using the second step of PCR using home-made indexing primers with completion of sequencing adapter sequences, then purified by SPRI beads and sequenced in paired-end mode using BGI service on DNBSEQ sequencing platform. We obtained about 30000 read pairs per transcript.

Sequencing data was aligned on expected transcript sequences using bowtie2 tool with default parameters. UMI sequences were extracted by umi_tools extract with bc-pattern=NNNNXNNNNXNNNN and then deduplicated by umi_tools dedup. SNP data was collected by bcftools mpileup with -a AD parameter, and then VCF files were parsed using home-mad Python script. For each animal, counts of C57BL/6 SNP were normalized by division on CAST SNP counts, resulting in a comparative level of allelic expression. Significance of these distributions differences were assessed by Mann-Whitney test and then filtered by Benjamini/Hochberg FDR correction with 0.1 p-value threshold.

### Data analysis

Hi-C data was analyzed using Juicer software [72] with slight modifications - particularly, we analyzed Hi-C data quality using custom DE and cis/trans metrics as described in at quality control steps described in [73]. ChIP-seq data analysis was performed according to ENCODE pipelines, as described in our previous works [74].

### Data availability

Sequencing data generated in this study are accessible via the NCBI BioProject PRJNA842410.

### Prediction changes in gene expression using Enformer model

We used the pytorch version of deep learning Enformer model from https://github.com/lucidrains/enformer-pytorch. We predicted gene expression using as input three DNA sequences with the center in TSS of three genes (*Cdh23, Slc29a3, Unc5b*). We run Enformer as for reference mm10 genome sequences as for DNA sequences with modeled mutations: 2 CTCF-bs (chr10:60,755,585-60,761,088 deletion) and 4 CTCF-bs (chr10:60,755,585-60,761,088 deletion, chr10:60,775,689-60,775,698 deletion and chr10:60,778,725-60,778,738 deletion with insertion of -AA- dinucleotide). We summed 4 bins around the TSS bin for predicted wild-type data and predicted mutated data, then we calculated the ratio between these two values to assess changes in gene expression for each gene. We used 5 target cell types from mouse Enformer target data: ‘liver, adult pregnant day01’, ‘urinary bladder, adult’, ‘kidney, neonate N30’, ‘cerebellum, adult’, and ‘olfactory brain, adult’.

## Author contributions

P.S. and V.F. conceived the studies and designed the experiment. P.S. and P.B. prepared gRNA and ssODN constructs; A.K. and I.S. performed microinjections; P.S. handled transgenic mice and performed genotyping with help from Y.S., A.Y., and S.T.; V.L and Y.S. performed ChiP-seq experiments; P.S., E.V. and P.B. performed NGS data analysis with help from V.F.; P.S. performed Hi-C experiments and N.T. analyzed the data; P.B. generated and analyzed Enformer predictions; P.S. prepared draft of the manuscripts with help from V.F.; all authors contributed to manuscript revision, read, and approved the submitted version.

## Conflict of Interest

The authors declare that the research was conducted in the absence of any commercial or financial relationships that could be construed as a potential conflict of interest.

## Acknowledgment

We acknowledge support of the RSF grants (2017-2019 #17-74-10143 and 2021-2024 #22-14-00247), which allowed us to build infrastructure for genetically-modified mice generation and NGS-based transcription analysis. The original manuscript text was composed by authors; proofreading was conducted with the assistance of ChatGPT o1. The text was corrected and edited by the authors after ChatGPT proofreading.

## Funding

This work was supported by the Ministry of Education and Science of the Russian Federation, agreement № 075-15-2024-539 (signed 24.04.2024).

## Notes

### Competing Interest Statement

The authors have declared no competing interest.

### Summary of Updates

Resolution of figures have been improved

