## Supplementary material 1 for "Direction and modality of transcription changes caused by TAD boundary disruption in *Slc29a3/Unc5b* locus depends on tissue-specific epigenetic context" (from another version of this preprint: version 2, 2024-12-26)

| Oligo ID | Sequence | Purpose |
| --- | --- | --- |
| T7+M1-GRNA_F | GTTAATACGACTCACTATAGTAAACTAAGACATCCAGTAGTTTTAGAGCTAGAAATAGCAAGTTAA | Components of CRISPR/Cas9 system for generation of [mm10] chr10:60755585-60761088 deletion |
| T7+R1-GRNA_F | GTTAATACGACTCACTATAGTAGAGAGAACAGGCGAACCTGTTTTAGAGCTAGAAATAGCAAGTTAA |  |
| DELM1R1_SSODN-TEMPLATE | TATTTGGGTCCTCACTTTCCAAGACTGTAGGTCTGCCTGCTCTCCCTTTTCTGCCCTTACTTCGCCTGTTCTCTCTACCCCACTCACCCAACGATAACCTCTCTCTCAAGGCAAGGACCC |  |
| T7+SGRNA_L2-CTCF | GTTAATACGACTCACTATAGTTTACAACTGCAGCCCCTGCGTTTTAGAGCTAGAAATAGCAAGTTAA | Components of CRISPR/Cas9 system for deleting of three telomeric CTCF-bs |
| T7+SGRNA_M2-CTCF | GTTAATACGACTCACTATAGTACCGAGAGCTCCCACCAGAGTTTTAGAGCTAGAAATAGCAAGTTAA |  |
| T7+SGRNA_R2-CTCF | GTTAATACGACTCACTATAGTGCCCCCTAGTGGCCACGGTGTTTTAGAGCTAGAAATAGCAAGTTAA |  |
| Unc5b_L2_SSODN-TEMPLATE | TGGAGAGGTGGTGAGAGGGATGGGAAAGAAAGA TGGAACCCAGGGAGCCCACAGCGAAAGCTTGCCAGGCAGGGGTCCACAGATCATTCTAGAAGGACAGAAATATCCTGTTCCCTTCAT |  |
| Unc5b_M2_SSODN-TEMPLATE | TCATCGTCTATGGTGTGGATGGGTGTTTGTG GCATGGGCTGTGATTATCCCGGGTGAAGCTTCTCTCGGTACTGTAGGGGGGCCTGAAGCAATATATCCCCTCCACTCTGCCCCACCCC |  |
| Unc5b_R2_SSODN-TEMPLATE | GCAAGTCTGAGAAGAGTTCCAAGAACATCAAGG GATTGAGCATGTGAGCATGAAATGAAGCTTCATGCCTATGGAACTGTTTACAGGAGCCCTTCACCTGCGGAAACAGGGTCCCCCTCC |  |
| REV | AAAAGCACCGACTCGGTGCCACTTTTTCAAG | Oligonucleotides for CRISPR/Cas9 sgRNA construction |
| SGRNA_REV | AAAAGCACCGACTCGGTGCCACTTTTTCAAGTTGATAACGGACTAGCCTTATTTTAACTTGCTATTTCTAGCTCTA |  |
| M1-F | GGAATTCAGGTTGAGTGGGC | Primers for [mm10] chr10:60755585-60761088 deletion genotyping |
| M1-R | GATCCCAGGTATAATGAACGC |  |
| R1-F | TCATCGCCGGGAAGTGTTAC |  |
| R1-R | CATCTCTCCAGCCCGGTAAAG |  |
| M2-F | CTGGGTACCTGGGATTTGGC | Primers for PCR-RFLP based telomeric CTCF-bs genotyping |
| M2-R | TCCTGCTCACCATTGACCAC |  |
| L2-F | ATTGGATGGGAGGTGCAGTG |  |
| L2-R | CGGTGCTTCTTAGCACTCCT |  |
| R2-F | CTCATGATCCAAGGGTACCTCC |  |
| R2-R | CTCCCACTGTATGCCAGACGA |  |
| R2-BARCODE-F | AC NNNN ACCTCATGATCCAAGGGTACCTCC | Primers for NGS-based telomeric CTCF-bs genotyping. NNNN - non-random predesigned sample-specific barcode |
| R2-BARCODE-R | AC NNNN AGCTGGAGGATTCCAGGCGA |  |
| M2-BARCODE-F | AC NNNN AGAGTGTGGCTGGGTACCTG |  |
| M2-BARCODE-R | AC NNNN AACTACACCTGTGTGGCCAAG |  |
| L2-BARCODE-F | AC NNNN AGCACGTGGGGTGAGAG |  |
| L2-BARCODE-R | AC NNNN CCACATTAGGTCTACTTGAGCA |  |
| Unc5b_ad_F3 | GGCTTCCACCCTGTCAACTTC | Primers and hydrolysing probes for Digital PCR |
| Unc5b_ad_R3 | GGCCAGACCCAGAACCGAT |  |
| Unc5b_ad_P3snp | HEX-CTGGAGTTATAGACCTTtATCTTGAGGCT-BHQ1 |  |
| Unc5b_ad_P3wt | FAM-CTGGAGTTATAGACCTTGATCTTGAGGCT-BHQ1 |  |
| Slc29a3 _ad_F1 | AATAGCAGCGTTTACGGCCT |  |
| Slc29a3 _ad_R2 | GACTGCTGCCATGAGGAAGAA |  |
| Slc29a3 _ad_P2wt | FAM-TGCTGTCCCGCACATCACTGGA-BHQ1 |  |
| Slc29a3 _ad_F2snp | HEX-TGCTGTCCCGCACGTCACTGGA-BHQ1 |  |
| Psap_ad_F1 | GAAAGATAATGCTACGCAGGAGGA |  |
| Psap_ad_R1 | TTGAGCGCAGAGCACACTTC |  |
| Psap_ad_P1wt | FAM-TCGTGCAAGGAGGTGGTTGACTCTT-BHQ1 |  |
| Psap_ad_P1snp | HEX-TCGTGTAAGGAGGCGGTTGACTCTT-BHQ1 |  |
| Cdh23_ad_F1 | CCAAGGTGGACTCTACTGTGAAG |  |
| Cdh23_ad_R1 | TCTCAAAGGCCCCAGCAATG |  |
| Cdh23_ad_P1wt | TAMRA-CGACTTCACCTCGGACTCAGCCATC-BHQ2 |  |
| Cdh23_ad_P1snp | ROX-CGACTTTACCTCGGACTCGGCCATC-BHQ2 |  |
| Vsir_ad_F1 | CCCTGCTCCTTGCTATTTTCCT |  |
| Vsir_ad_R1 | ATCTGGACCTCGCCTCGT |  |
| Vsir_ad_P1wt | TAMRA-TCTTGTAGATGGTCACATCGTGCCCTT-BHQ2 |  |
| Vsir_ad_P1snp | ROX-TCTTGTAGACGGTCACATCGTGCCCTT-BHQ2 |  |
| Sgpl1_ad_F | TGTGAAAACTCTGCCTGCTCA |  |
| Sgpl1_ad_R | ATATCTGGATGCAGTGGATTGCT |  |
| Sgpl1_ad_Pwt | TAMRA-AGCTGTGTACAATGGGGAACCGAAGCTCA-BHQ2 |  |
| Sgpl1_ad_Psnp | ROX-AGCTGTGTACAACGGGGAACCGAAGCTCA-BHQ2 |  |
| ActB_F | TATTGGCAACGAGCGGTTCC |  |
| ActB_R | TGGCATAGAGGTCTTTACGG |  |
| ActB_P | Cy5-CCAGCCTTCCTTCTTGGGTATGGAAT-BHQ2 |  |
| adapterR | CTGGAGTTCAGACGTGT | Primers used for allele-specific NGS-based expression measurement |
| Slc29a3 -scstr1 | TCTTTCCCTACACGACGCTCTTCCGATCTTATCAGGAGGAGCCATGGGAG |  |
| Slc29a3 -rev1 | CTGGAGTTCAGACGTGTGCTCTTCCGATCTNNNNANNNNCNNNNCATACAGAGCCCAAGGAAGACT |  |
| Unc5b_2-scstr1 | TCTTTCCCTACACGACGCTCTTCCGATCTGAACAAGCCAGTGGAACTGC |  |
| Unc5b_2-rev1 | CTGGAGTTCAGACGTGTGCTCTTCCGATCTNNNNANNNNCNNNNACACCTCGATCTGCACCTCT |  |
| Vsir-2-scstr1 | TCTTTCCCTACACGACGCTCTTCCGATCTCTGGGGACGTCTTTTTCCCA |  |
| Vsir-2-rev1 | CTGGAGTTCAGACGTGTGCTCTTCCGATCTNNNNANNNNCNNNNATAGGGGAGGGTCTGTGCAT |  |
| Cdh23-1-dd-scstr1 | TCTTTCCCTACACGACGCTCTTCCGATCTACATTGACAGTCGTGGCAGA |  |
| Cdh23-1-dd-rev1 | CTGGAGTTCAGACGTGTGCTCTTCCGATCTNNNNANNNNCNNNNGCCTTGACTGTGGCTACTCG |  |
| Sgpl-scstr1 | TCTTTCCCTACACGACGCTCTTCCGATCTACAACAGGTGAGCAAAGCCA |  |
| Sgpl-rev1 | CTGGAGTTCAGACGTGTGCTCTTCCGATCTNNNNANNNNCNNNNTTGCTCCACGTGAATTCTCCA |  |
| Psap-scstr1 | TCTTTCCCTACACGACGCTCTTCCGATCTTCACCGAAGCTGGGAACTTG |  |
| Psap-rev1 | CTGGAGTTCAGACGTGTGCTCTTCCGATCTNNNNANNNNCNNNNTCCAGGATGACAGGCAGGTA |  |
| Supplementary Table 1. Oligonucleotides used in work | | |

| Mouse | Centromeric deletion | Left | | Middle | | Right | |
| --- | --- | --- | --- | --- | --- | --- | --- |
|  |  | RFLP | NGS | RFLP | NGS | RFLP | NGS |
| 1 | + |  |  | + | + |  |  |
| 2 |  |  |  | + | + |  |  |
| 3 | + |  |  | + | + |  |  |
| 4 | + |  |  |  | + | + | + |
| 5 |  |  |  |  | + |  |  |
| 6 | + |  |  | + | + |  |  |
| 7 | + |  |  | + | + |  |  |
| 8 |  |  | + | + | + |  | + |
| 9 |  | + | + | + | + |  |  |
| 10 |  |  | + | + | + |  |  |
| 11 | + | + | + |  | + |  |  |
| 12 |  | + | + | + | + |  |  |
| 13 | + | + | + |  | + |  | + |
| 14 | + |  |  | + | + |  | + |
| 15 | + | + | + |  | + |  | + |
| 16 | + | + | + | + | + |  |  |
| 17 | + |  |  |  |  |  |  |
| 18 | + |  |  | + | + |  |  |
| 19 | + |  |  |  |  |  |  |
| Supplementary Table 2. Genotyping of multiple CTCF-bs knock-out mice | | | | | | | |
